# *De novo* design of inducibly assembling multi-component filaments

**DOI:** 10.1101/2025.03.26.645066

**Authors:** Hao Shen, Eric M. Lynch, Noel Jameson, Justin Decarreau, Chenyang Shi, Zibo Chen, William Sheffler, Miaomiao Xu, James J De Yoreo, Jesse G. Zalatan, Justin M. Kollman, David Baker

## Abstract

The design of multi-component nanomaterials is an outstanding challenge. Here, we describe the computational design of protein filaments with two or three distinct structural components that assemble into micron-scale, well-ordered fibers when mixed. CryoEM structure determination of four fiber designs was close to the computational design models. Filament assembly can be initiated by mixing the components, and modulated by addition and/or phosphorylation of designed regulatory subunits. This work demonstrates that regulatable multi-component protein filament systems can now be designed, opening the door to a wide range of engineered materials.

## Introduction

Many natural protein filaments, such as microtubules^1,2^, are composed of two or more structural components, which allows for intricate regulation of their assembly, dynamics, and interactions within the cell. In contrast, *de novo*-designed protein filaments have predominantly been constructed from single protein components^3,4^. This approach, while successful in creating self-assembling structures, limits the complexity and functional versatility achievable in synthetic filaments.

We reasoned that the design of protein filaments with multiple structural components could enable multiple levels of control over filament assembly, stability, and functionality. We set out to design such systems, along with additional components that modulate their assembly, starting from *de novo*-designed heterooligomeric systems. We chose three such starting systems: (1) designed helical hairpin heterodimers (DHDs)^5^ and (2) designed heterotrimers (DHTs)^6^, which both have interaction specificity determined by networks of buried hydrogen bonds, and (3) reversibly assembling beta-strand pairing-based heterodimers (LHDs)^7^. The advantage of the DHDs and DHTs is that by incorporating different buried hydrogen bond networks, very high interaction specificity has been achieved, enabling, in principle, the design of many orthogonal interacting filament systems. The advantage of the LHDs is that the components are well-behaved proteins in isolation since much of the heterodimer interface is a polar edge beta-strand. We describe each of these three in turn in the following sections.

## Results

### Fiber design and characterization

DHDs and DHTs are designed heterodimers and heterotrimers in which the monomeric subunits are helical hairpins, and the chains interact via buried hydrogen bond networks that confer specificity. In the DHD case, 21 distinct pairs have been designed; in all-by-all mixing experiments, each subunit pairs specifically with its partner^5^. This multiplicity and orthogonality make the DHDs and DHTs attractive building blocks for higher-order materials.

We sought to construct DHD-based filaments using the design models of DHD131, DHD37_1:234, DHD127, and DHD15^5^. We used the Rosetta-based helical docking and design method described in Shen *et al.*^3^ to design interfaces on these building blocks predicted to direct assembly into protein filaments. To ensure that fiber assembly requires each heterodimer subunit (such that the monomers alone cannot assemble), we required that the fiber interfaces had contributions from each subunit (Fig. 1A). We generated 195,000 helical filament backbones and selected 55 designs for experimental testing [we refer to these below as DHD_HFs (DHD-based two-component helical filaments)] based on the selection criteria described in the Methods section. For the DHT-based filaments, we used the crystal structure of heterotrimer DHT03^6^ as the starting scaffold, generated 163,900 helical filament backbones (Fig. 1B, we again required that the filament interfaces involve all subunits), and selected 41 designs for experimental testing [we refer to these below as DHT_HFs (DHT-based three-component helical filaments)].

**Fig. 1.**
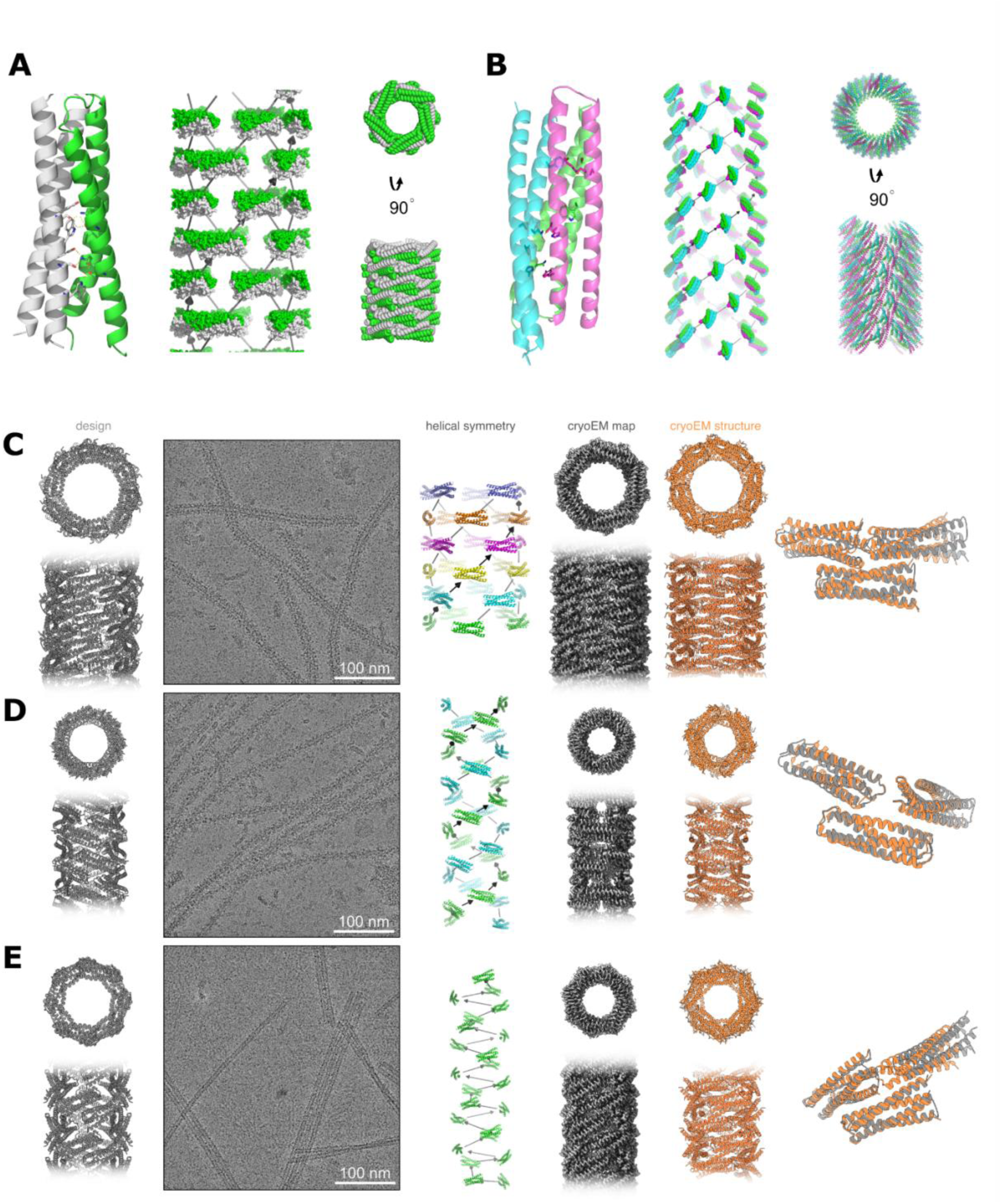
Design and structure of multi-component helical bundle protein filaments. (**A**) (left) Two-component heterodimer DHD131 as the scaffold for filament generation. Computational sampling of alternative helical packing arrangements of subunit shown in (middle), with each component colored. The grey arrows represent the rigid body transform that generates the helical assembly. (right) Compact packing arrangement of fibers subunits in lowest energy assemblies following sequence design. (**B**) (left) Designed heterotrimer DHT03 as the scaffold for filament generation. Computational sampling of alternative helical packing arrangements of subunit shown in (middle), with each component colored. The grey arrows represent the rigid body transform that generates the helical assembly. (right) Compact packing arrangement of fiber subunits in lowest energy assemblies following sequence design. (**C**-**E**) CroyEM structures of DHD-based two-component filaments. Computational model (first panel), representative filaments in cryoEM micrographs (second panel), helical symmetry (third panel), cryoEM map (fourth panel), cryoEM structure (fifth panel) and overlay between model and structure (sixth panel) for (**C**) DHD_HF30 - r.m.s.d. 1.4 Å, (**D**) DHD_HF20 - r.m.s.d. 2.0 Å, and (**E**) DHD_HF3 - r.m.s.d. 6.0 Å.

LHDs are designed heterodimers with interfaces containing an extended beta-sheet spanning the two subunits. The exposed beta strands at the subunit interfaces (prior to complex formation) have multiple polar groups and, hence, are soluble and well-behaved when monomeric. While there are fewer distinct LHDs than DHD heterodimers, the former have the advantage of reversible assembly and disassembly. We chose LHD317^7^ for LHD-based filament as it has the most alpha-helical surface for docking, generated 74051 helical filament backbones (Fig. 2A,B), and selected 70 designs for experimental testing [we refer to these as LHD-based filament (LHD_HFs)].

**Fig. 2.**
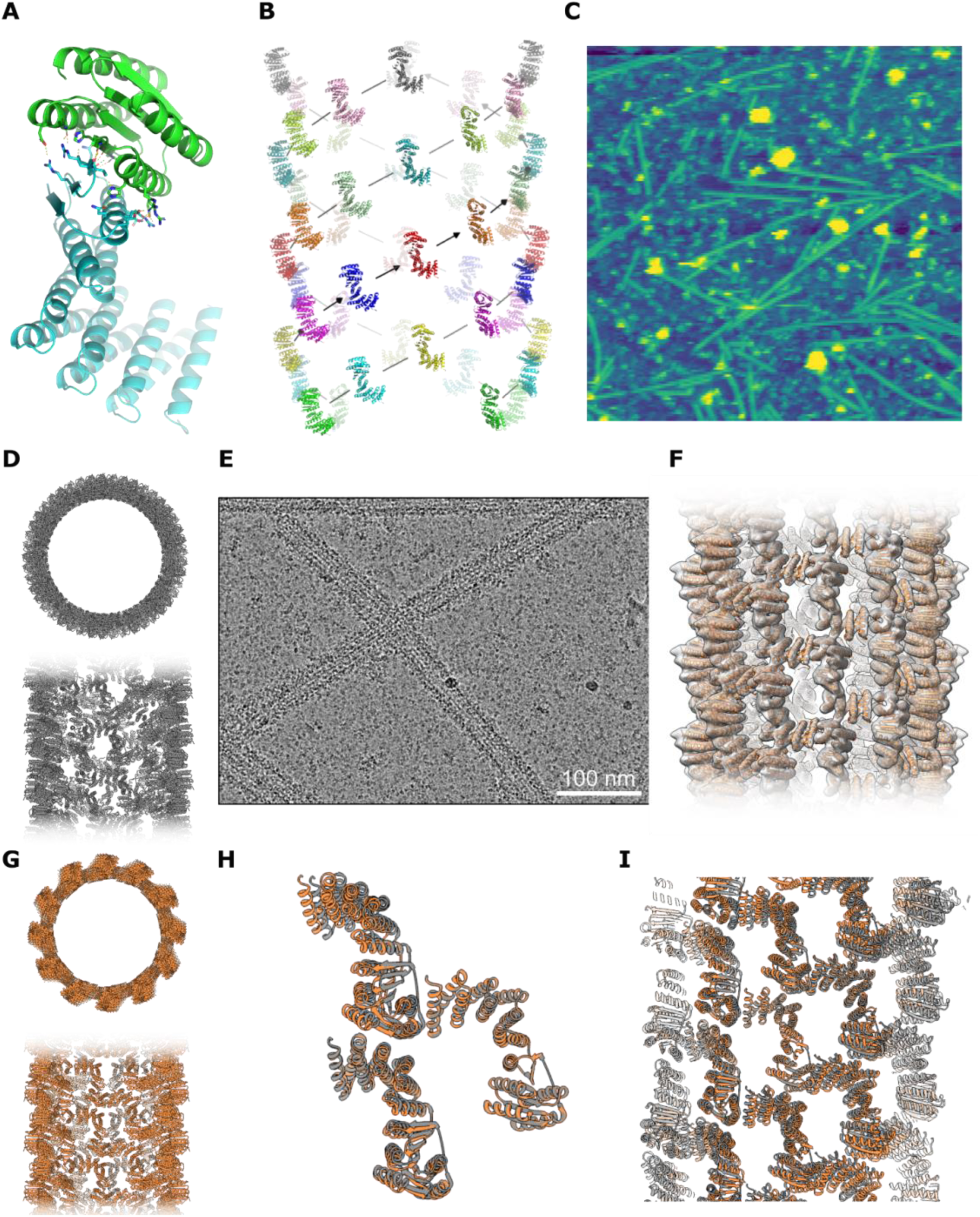
Design and structure of multi-component protein filaments built from reconfigurable LHD heterodimers. (**A**) The reconfigurable heterodimer LHD317 scaffold. (**B**) Helical symmetry of LHD_HF59 fiber; subunits related by C4 symmetry have the same color. (**C**) AFM micrograph of LHD_HF59. (**D**) Top and side view of LHD_HF59 design model (**E**) CryoEM micrograph of LHD_HF59. (**F**) LHD_HF59 filament structure fits into the cryoEM map. (**G**) Top and side view for LHD_HF59 structure. (**H,I**) Overlay between the design model and cryoEM structure at the designed interface (**H**) and in the overall helical architecture (**I**).

The components of the selected DHD_HFs, DHT_HFs, and LHD_HFs were co-expressed in *Escherichia coli* using a bicistronic or tricistronic plasmid under the control of a T7 promoter and purified by immobilized metal affinity chromatography (IMAC, only one of the components contains hexahistidine tag). The non-histagged component for 45 of the 55 selected DHD_HFs and for all of the selected LHD_HFs were fused to a green fluorescent protein (GFP) to facilitate screening and kinetic measurements. For the remaining 10 DHD_HFs, there was insufficient space to accommodate a GFP fusion without disrupting the design; their expression was confirmed by SDS-PAGE, and fiber formation was assessed by negative-stain electron microscopy. For the 70 three-component DHT_HFs, GFP was fused to only one of the non-his-tagged subunits.

Of the 55 selected DHD_HFs designs, 43 were well-expressed and co-recovered in the IMAC eluate. Almost all of the DHT_HF and LHD_HF components were also expressed and recovered. For all three cases, IMAC eluates were concentrated, and filament formation was monitored by electron microscopy (EM) and fluorescence microscopy. A total of six DHD_HF designs were found to form 1D nanostructures over 200 nm by EM (Fig. S1). One of each of the DHT and LHD designs, DHT_HF2 and LHD_HF59, were found to form filaments by negative stain EM (nsEM) and fluorescence methods (Fig. S2). LHD_HF59 filaments with uniform diameters were also observed with atomic force microscopy (AFM) (Fig. 2C). The presence of each co-expressed component was confirmed by SDS-PAGE (Fig. S3).

We chose three DHD_HF designs and LHD_HF59 with a range of model architectures and highly ordered nsEM morphologies for higher resolution structure determination by cryo-electron microscopy (cryoEM). We determined the filament structures and refined helical symmetry parameters using iterative helical real space reconstruction in SPIDER^8,9^ or Relion^10,11^, followed by further 3D refinement in cryoSPARC^12^. The DHD_HF structures were solved to 3.2-3.3 Å resolution and LHD_HF59 to 5.9 Å. In all four cases, the overall orientation and packing of the monomers in the filament were similar in the experimentally determined structures and design models, but there was considerable variation in the accuracy with which the details of the interacting interfaces were modeled, driven largely by differences observed in DHD_HF3 (see below).

Three of the four designed filaments matched the computational models at near-atomic resolution. For DHD_HF20 and DHD_HF30, the experimentally observed rigid body orientations were nearly identical to the design models (1.4 Å and 2.0 Å r.m.s.d. over three chains containing all unique interfaces, respectively). For both structures, the backbone and side chain conformations at the subunit interfaces were very similar to those in the design model. In contrast, the DHD_HF3 structure exhibited larger deviation from the computational design: the r.m.s.d over three interface chains was 6Å, while the primary and secondary interfaces had r.m.s.d values of 1Å and 5.5Å, respectively. Higher resolution cryoEM maps showed that the main interface deviation was due to the backbone deviation where the design model and crystal structure of the scaffold DHD131 disagree (Fig. S4)—as our computational method assumes a rigid backbone, building block backbone deviation leads to deviation at the interface level. The helical symmetry of DHD_HF30 has a rise of 22.3 Å, rotation of 37.1°, and a larger diameter of 11.6 nm, with additional C5 point group symmetry along the helical axis. The structure can be viewed as a stacked C5 fiber with helical rotation, with the helices in the subunits perpendicular to the helical axis (Fig. 1C). DHD_HF20 has a rise of 15.2 Å, rotation of 66.5° and diameter of 8.7 nm, with additional C2 point group symmetry along the helical axis; in cross-section six subunits form a ring (Fig. 1D). DHD_HF3 has a rise of 5.9 Å, rotation of −135.9° (left-handed helix), and diameter of 9.9 nm; in cross-section five subunits form a ring (Fig. 1E).

For LHD_HF59, the 5.9 Å cryoEM structure has 1.94 Å r.m.s.d. to the design model over three chains (Fig. 2D-I). The helical symmetry of LHD_HF59 relating non-contacting subunits has a rise of 23.5 Å, rotation of 119.3°, and a large diameter of 23.5 nm, with additional C4 point group symmetry along the helical axis (Fig. 2B). The resulting overall architecture can be viewed as a right-handed helix composed of four identical, parallel strands with non-contacting subunits (or left-handed helix composed of eight identical, parallel strands with contacting subunits), with the beta-sheet interface within the LHD is perpendicular to the fiber axis. The close agreement between the design model and cryoEM structure for three of the four designs shows that computational methods can now accurately design multicomponent fibers.

An attractive feature of multicomponent fibers is that assembly should be triggerable by mixing. To investigate this, we expressed and purified the components of the DHD_HF30 fiber separately, and then mixed them to initiate fiber assembly. NsEM showed that filaments formed after mixing (Fig. 3A-D), this was verified quantitatively using fluorescent microscopy (fiber count refers to the number of fiber-like objects identified based on automated segmentation; see Methods).

**Fig. 3.**
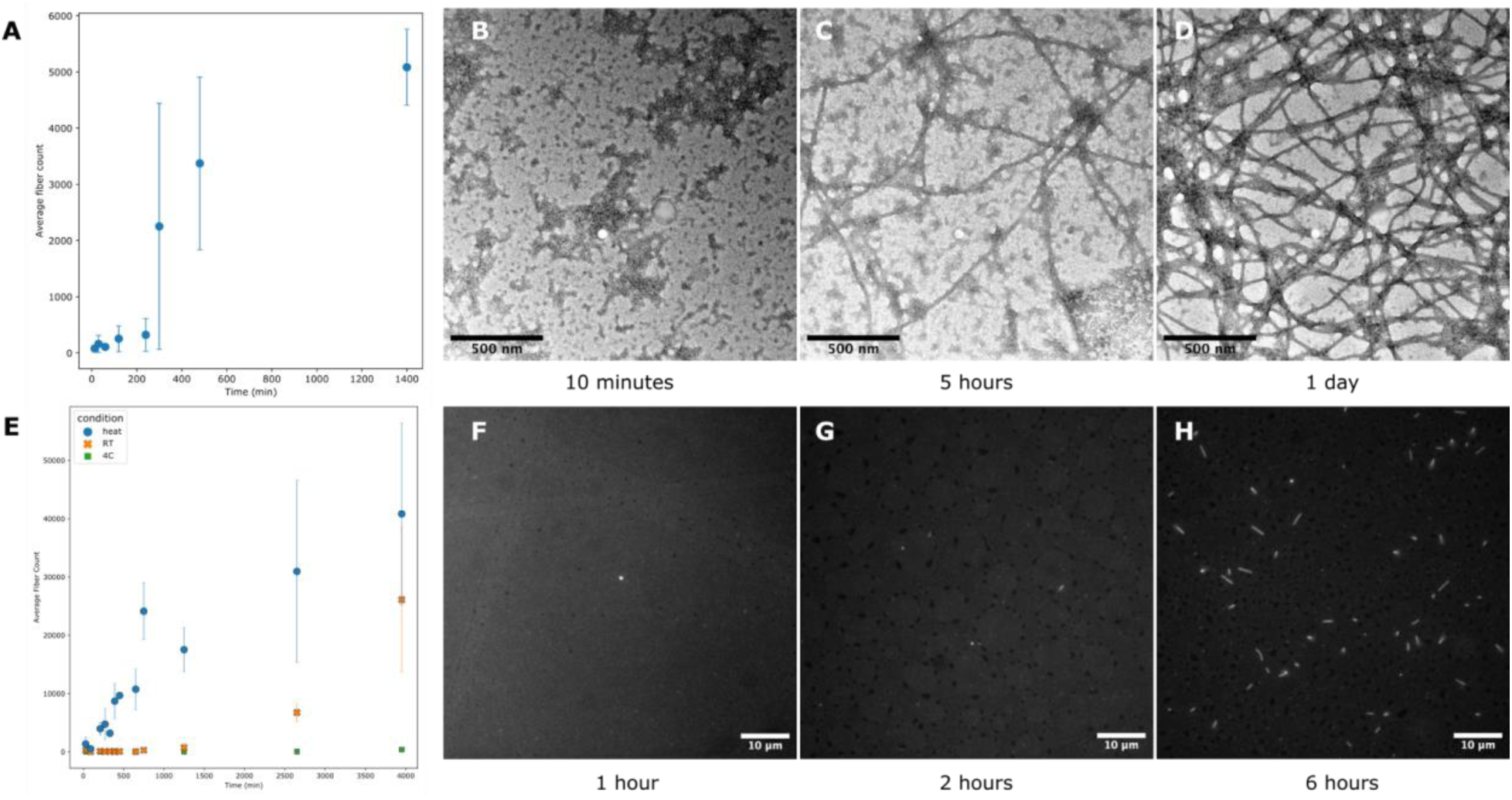
Fiber formation by mixing components. (**A**) Fluorescent microscopy quantification of fiber count ± SEM at different times after mixing components for DHD_HF30. (**B-D**), DHD_HF30 assembly after 10 minutes (**B**), 5 hours (**C**), and 1 day (**D**), characterized by negative stain electron micrograph. (**E**) Fluorescent microscopy quantification of fiber count for LHD_HF59 fiber growth upon mixing over time at different temperatures. Heat refers to 60°C incubation for the first hour, and room temperature afterwards. RT refers to room temperature all the time. 4C refers to incubation at 4°C. **(F-H)** Representative fluorescent microscopy images of LHD_HF59 after 1 hour at 60°C (**F**), then incubated at room temperature for 2 hours (**G**), and 6 hours (**H**), characterized by fluorescent microscopy.

We took advantage of the fact that the LHD fiber building blocks, in addition to being well-behaved in isolation^7^, are larger and hence have termini spaced further apart to fuse both components to fluorescent proteins. We began by characterizing the growth dynamics of LHD_HF59, in which both components are fused to GFP. We found that temperature affects both nucleation and fiber growth rates. Heating the protein mixture to 60°C for one hour—although fibers do not form during this step—resulted in faster fiber formation over the subsequent hours (Fig. 3E–H), compared to slower formation overnight at room temperature and no fiber formation after a month at 4°C. LHD_HF59 fibers are long enough to allow quantification of average fiber length over time (Fig. S5A), which is difficult for other fibers around 1 μm due to the resolution limits of fluorescence microscopy.

We found that fusing different fluorescent protein fusions to component B affect the fiber formation rate. Fusing component B to mScarlet-I, a red fluorescent protein, significantly slowed fiber formation, taking up to five days (Fig. S5B). A point mutation at the fiber interface of component B—from the polar residue lysine to the nonpolar residue leucine—accelerated assembly when component B was fused to mScarlet-I. In this case, fibers formed within 24 hours (Fig. S5C), likely due to increased hydrophobic interactions promoting assembly. Storing the component B in buffers containing 400 mM imidazole also accelerated fiber formation to within hours (Fig. S5D).

To regulate the assembly process for DHD_HFs, we designed shorter complementary pairing subunits without fiber interfaces (called “blockers”) to coexpress with the individual components (Fig. S6 A,B). Defining the original heterodimer as A:B, and the redesigned fiber-forming version as A0:B0, we experimented with expressing A0 with a shorter version of B (called “b”), and B0 with a shorter version of A (called “a”). We hypothesized that when the A0:b complex is mixed with a:B0 complex, fiber assembly would be promoted as the full-length A0:B0 complex has the most extensive interface and the lowest free energy. To tune the assembly rate, the blockers were truncated to different lengths. This blocker strategy also alleviates the tendency of the DHD subunits to form homodimers when expressed alone, which would otherwise hinder filament formation.

We tested this assembly regulation strategy with DHD_HF30. Fiber component A0 was co-expressed with blocker b, fiber component B0 was co-expressed with blocker a, and the complexes were purified. Both blockers have ∼7 residues (one heptad) truncations at each terminus. When the blocker A0 and B0 complexes were mixed, fibers formed as hypothesized. The fiber formation rate increased with higher temperatures (Fig. S6 C,D), which likely helps overcome the energy barriers to the dissociation of the blocked complexes.

Regulation of fiber assembly by phosphorylation could enable the design of signaling pathway-responsive extensions to the cytoskeleton for synthetic biology. To explore the possibility of kinase-regulated assembly, we incorporated PKA kinase recognition sequences^13^ into the heterodimeric interface of full-length blockers. We anticipated that phosphorylation of these sites would disrupt the blocked complexes, which would release assembly-competent monomers that could then assemble (Fig. 4A). We found by both nsEM and fluorescent microscopy that adding PKA to such blocked assemblies at 37 °C resulted in substantial stimulation of fiber assembly compared to control without kinase (Fig. 4B-E).

**Fig. 4.**
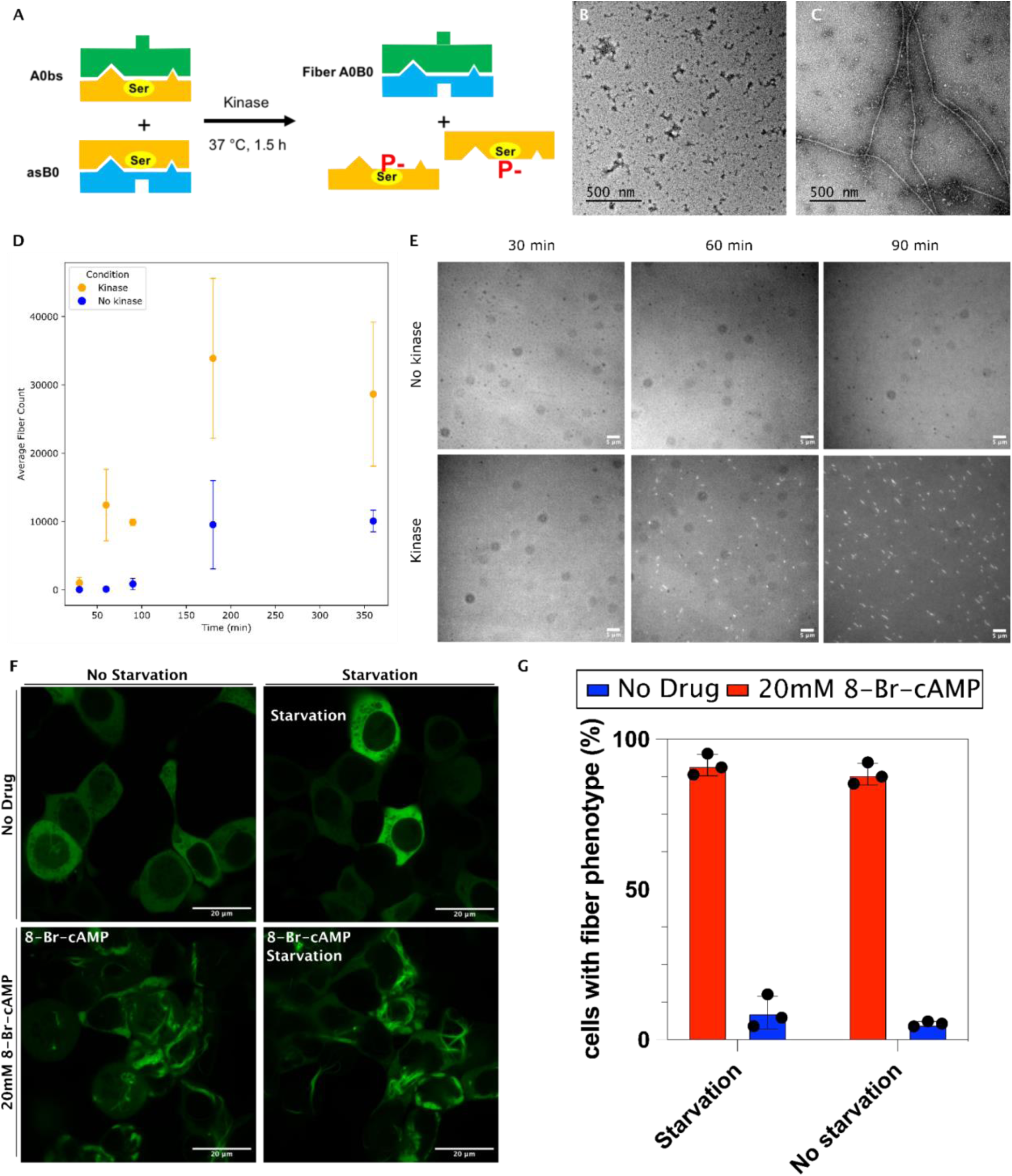
Kinase-dependent fiber assembly. (**A**) Cartoon representation of the induced assembly scheme. Components are purified with blockers that are released through phosphorylation by PKA. **(B,C)** Assembly characterized by negative stain EM in the (**B**) absence and (**C**) presence of PKA incubated at 37°C for 1.5 hours. (**D**) Quantification by fluorescent microscopy of fiber number count (± SEM) and (**E**) Representative fluorescent microscopy images (**F**) Super-resolution confocal spinning disk microscopy images of fibers in HEK293T cells without (left panels) and with (right panels) 8-Br-cAMP. Cells were grown either in 10% FBS (panels 1 and 3) or 0% FBS (panels 2 and 4). Long fibers were observed following kinase induction by drug under both growth conditions. (**G**) Quantification of the percent of cells displaying the fiber formation phenotype in uncropped Z-stack images in triplicate. Statistical significance was determined with an unpaired T-test between cells dosed with 20mM 8-Br-cAMP and DMSO. For serum starved cells, P value = 2.6*10^-5^, for non-starved cells, P value = 3*10^-6^.

We proceeded to test whether kinase-regulated assembly could occur in mammalian cells. We transiently transfected plasmids encoding both fiber proteins and their associated regulatory subunits into HEK293T cells. To induce fiber formation, we subjected the cells to 8-Br-cAMP, a PKA activator. Twenty-four hours after drug administration, cells were imaged using spinning disc confocal microscopy to assess fiber formation (Fig. 4F). In the absence of the drug, there is a low basal level of fiber formation, with fibers observed in ∼10% of cells. There was a substantial and statistically significant increase in fiber formation in the presence of 8-Br-cAMP to ∼95% of cells (Fig. 4G), indicating that *in vivo* fiber formation can be modulated by PKA phosphorylation. Fiber levels were the same when cells were grown in starvation (0% FBS) or no starvation (10% FBS) conditions, indicating that there was no background activation due to PKA-stimulating ligands in FBS media.

## Discussion

This work demonstrates the *de novo* design of multi-component protein filaments that assemble into well-ordered, micron-scale fibers. Having multiple structural components, like tubulin (composed of a- and b-tubulin), provides greater functional versatility and multiple routes to dynamic regulation compared to previously designed single-component fibers. Filaments assemble only upon mixing components, demonstrating conditional assembly akin to natural systems. Assembly can be further regulated using blocker components, which can be released following phosphorylation by kinases in cells, opening avenues for constructing responsive materials and integrating synthetic fibers into signaling pathways. LHD_HF59, with its hollow interior, could enable molecular encapsulation, diffusion-based transport, and long-range signaling. Overall, this work establishes a versatile framework for designing programmable, multi-component protein filaments with dynamic regulation and modular functionality, which could enable applications in biosensing, drug delivery, bioelectronics, and synthetic biology, and pave the way for next-generation engineered materials.

## Acknowledgments

We thank S. Gerben, A. Murray, P. Heine, M. DeWitt, and R. Ravichandran, for their help with protein production. We also thank the Arnold and Mabel Beckman Cryo-EM Center at the University of Washington for the use of electron microscopes. **Funding:** This work has been supported by the Howard Hughes Medical Institute (H.S., W.S., and D.B.), The Audacious Project at the Institute for Protein Design (H.S. and D.B.), Bill & Melinda Gates Foundation grant INV-043758 (J.D.), Defense Advanced Research Projects Agency Biostasis HR001118S0034 (H.S.), NIH R35GM124773 (J.G.Z, N.J.), Department of Energy grant DE-SC0019288 and Alfred P Sloan Foundation G-2021-16899 (E.M.L.), NIH grants R35GM149542, S10OD032290 (J.M.K.), and 5P41GM103533 (Z.C.), and the Center for the Science of Synthesis Across Scales (CSSAS) as part of the US DOE BES Energy Frontier Research Center program under Award DE-SC0019288 at UW and FWP 77248 at PNNL. **Author contributions:** H.S. and D.B. designed the project. H.S. carried out the design calculations. M.X. assisted in the design selection. H.S. expressed, purified, and screened design proteins by negative stain EM. E.M.L., H.S., and J.M.K. carried out the cryo-EM data acquisition and structure determination. H.S. and J.D. performed fluorescence microscopy for fiber in solution and quantification. N.J. and J.G.Z. carried out *in vivo* phosphorylation assay by fluorescence microscopy and quantification. C.S. and J.J.D.Y. carried out the AFM data acquisition and analysis. H.S., W.S., and D.B. designed the method figure. H.S. and D.B. wrote the manuscript.

---

**Fig. S1.**
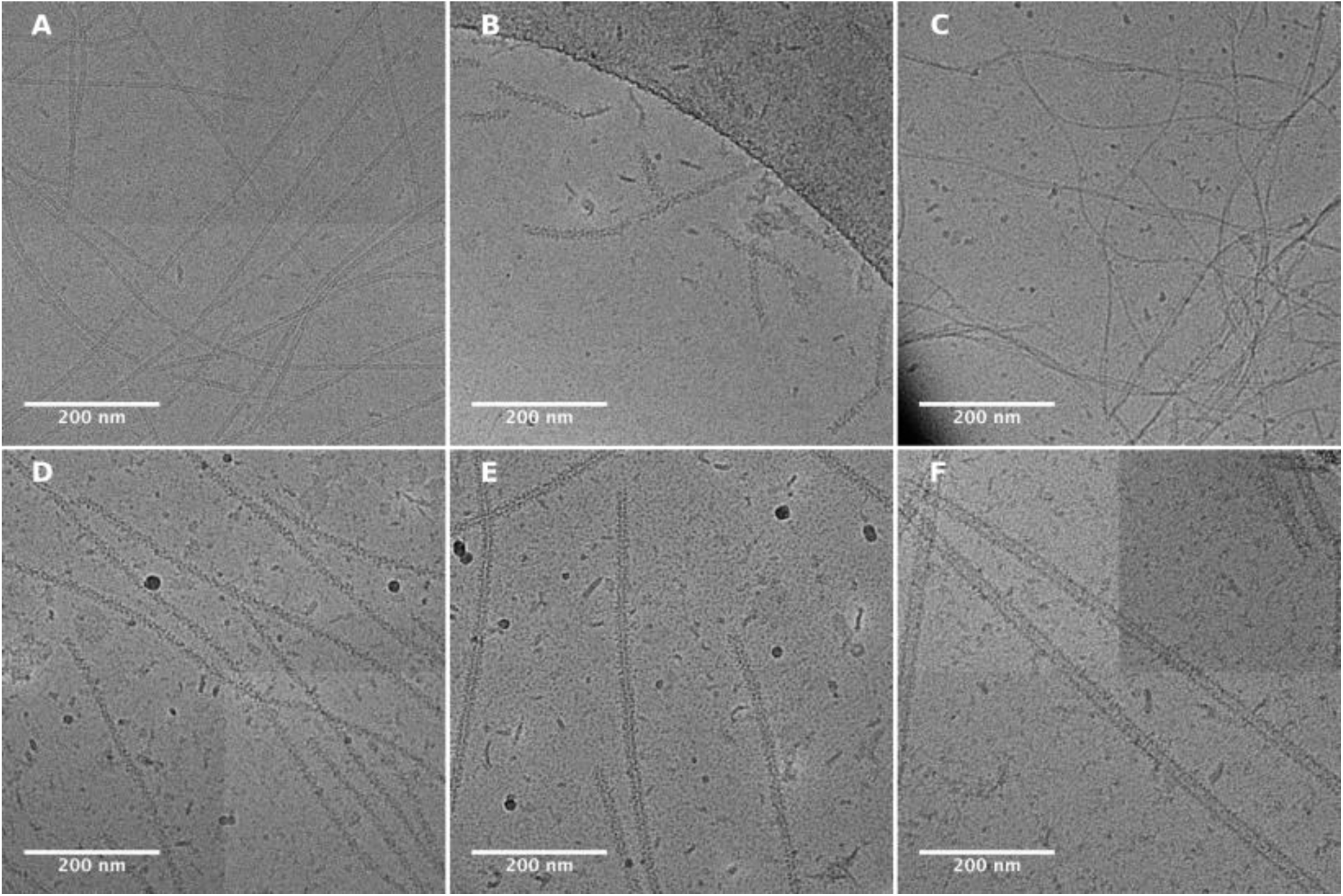
Cryo-electron micrograph of two-component filaments. (**A**) DHD_HF3. (**B**) DHD_HF16. (**C**) DHD_HF18. (**D**) DHD_HF20. (**E**) DHD_HF30. (**F**) DHD_HF41.

**Fig. S2.**
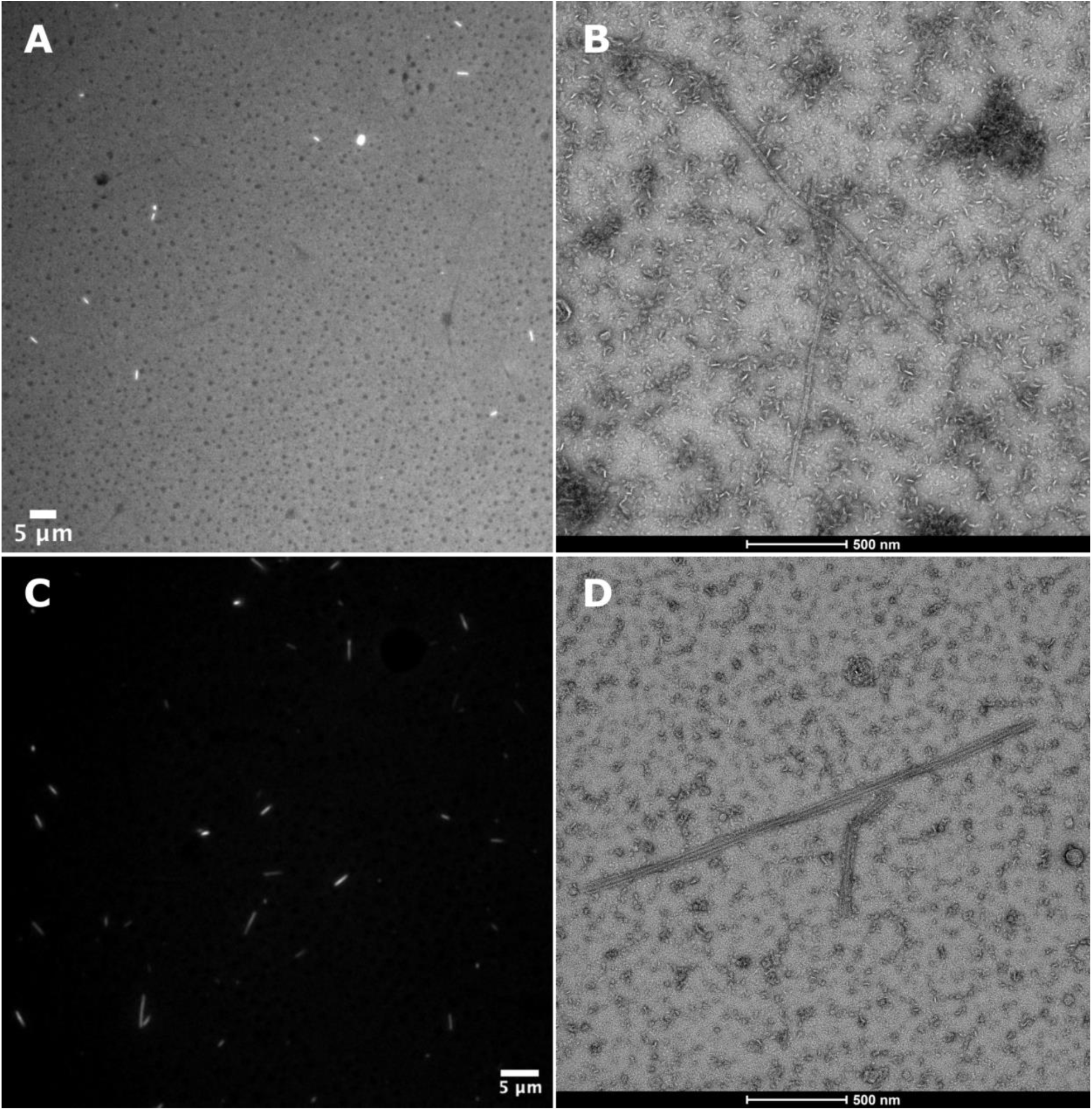
Fluorescent micrograph and negative stain electron micrograph of DHT_HF2 and LHD_HF59. (**A**) DHT_HF2 Fluorescent micrograph and (**B**) DHT_HF2 negative stain EM. (**C**) LHD_HF59 Fluorescent micrograph and (**D**) LHD_HF59 negative stain EM.t

**Fig. S3.**
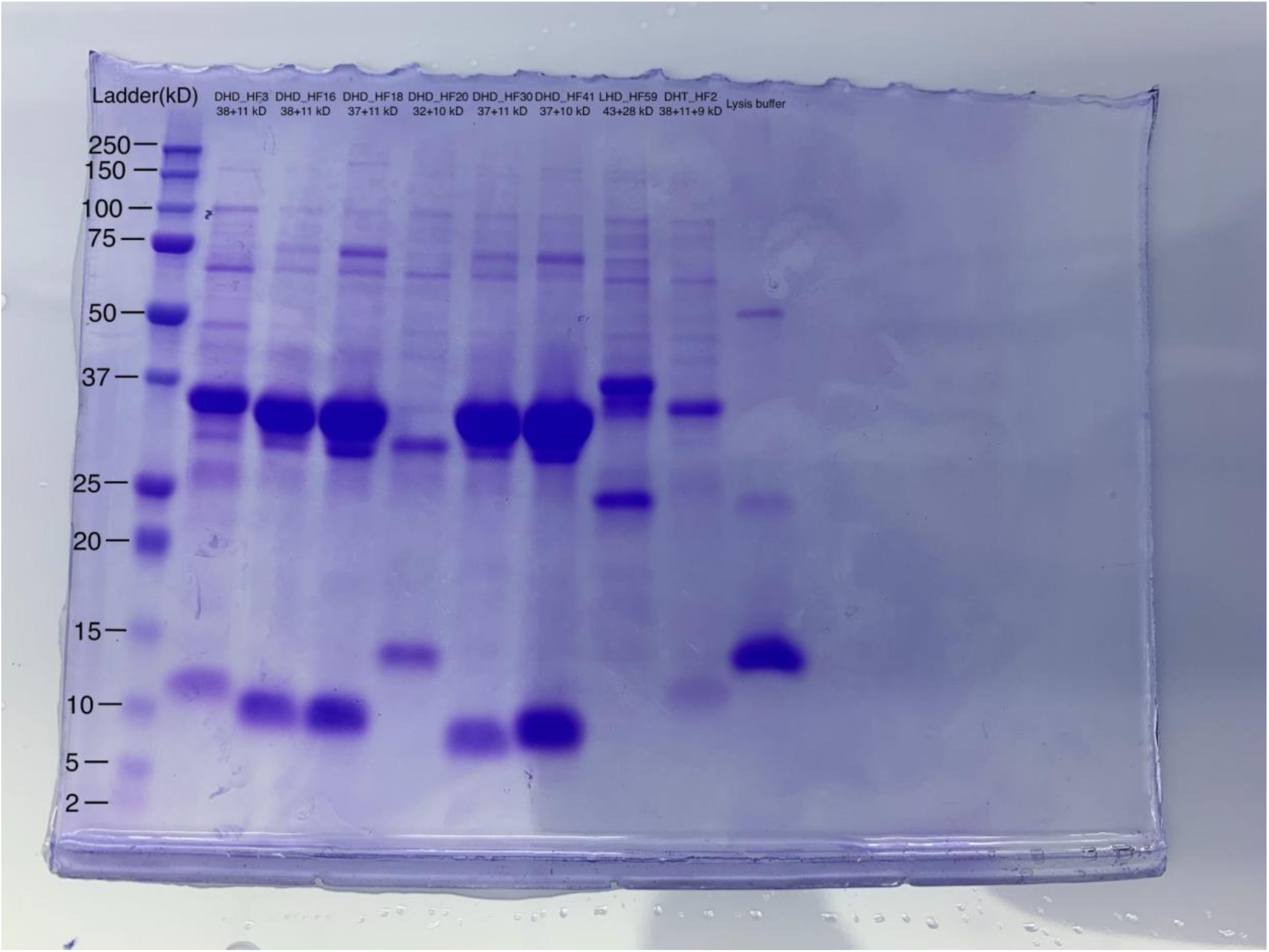
SDS-PAGE of DHD_HF3, DHD_HF16, DHD_HF18, DHD_HF20, DHD_HF30, DHD_HF41, LHD_HF59, and DHT_HF2 co-expression, shown in lane order.

**Fig. S4.**
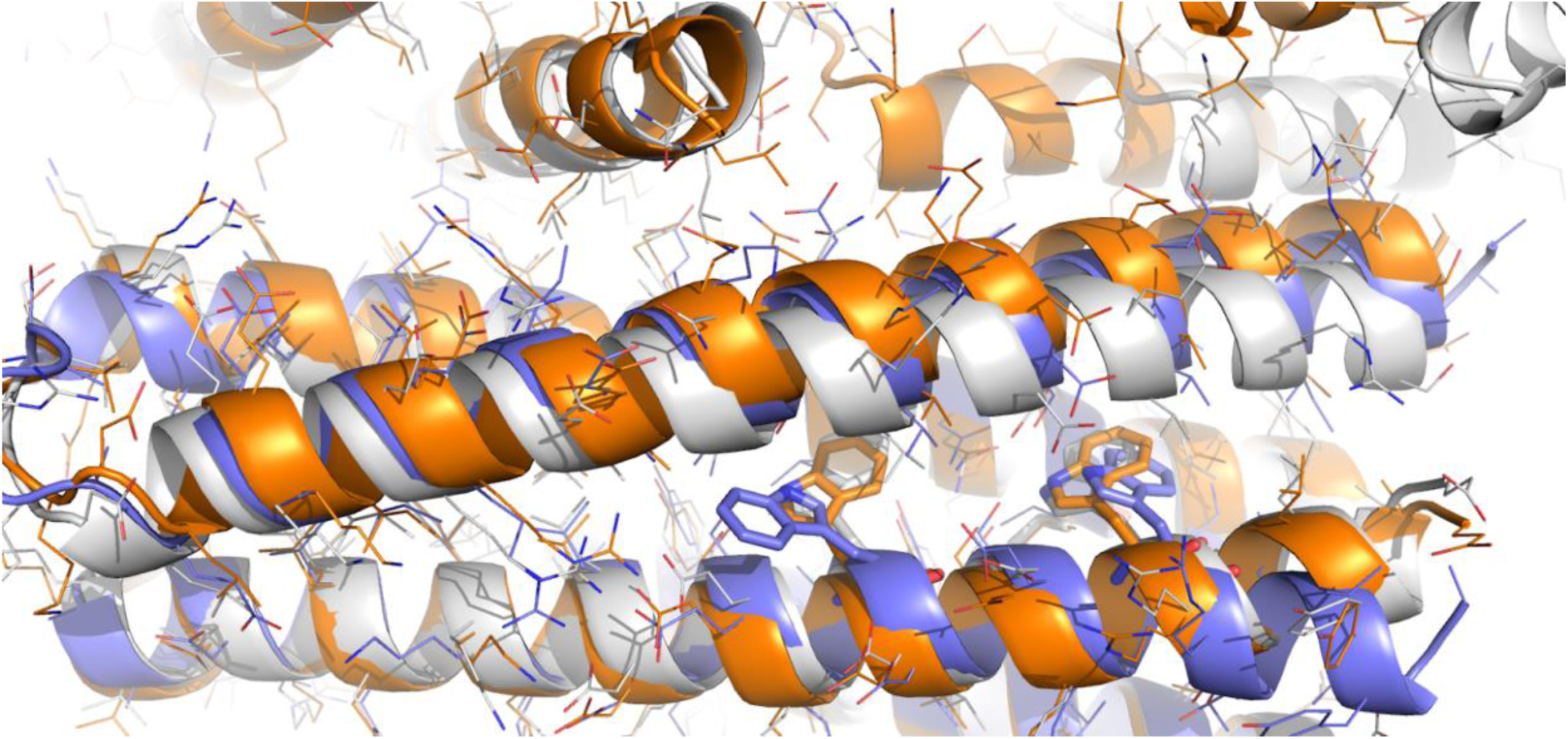
Comparison of the deviated interface for DHD_HF3. Aligned on one subunit are the DHD_HF3 design model (grey), cryoEM structure (orange), and the crystal structure for its scaffold DHD131 (purple) (PDB id:6DKM).

**Fig. S5.**
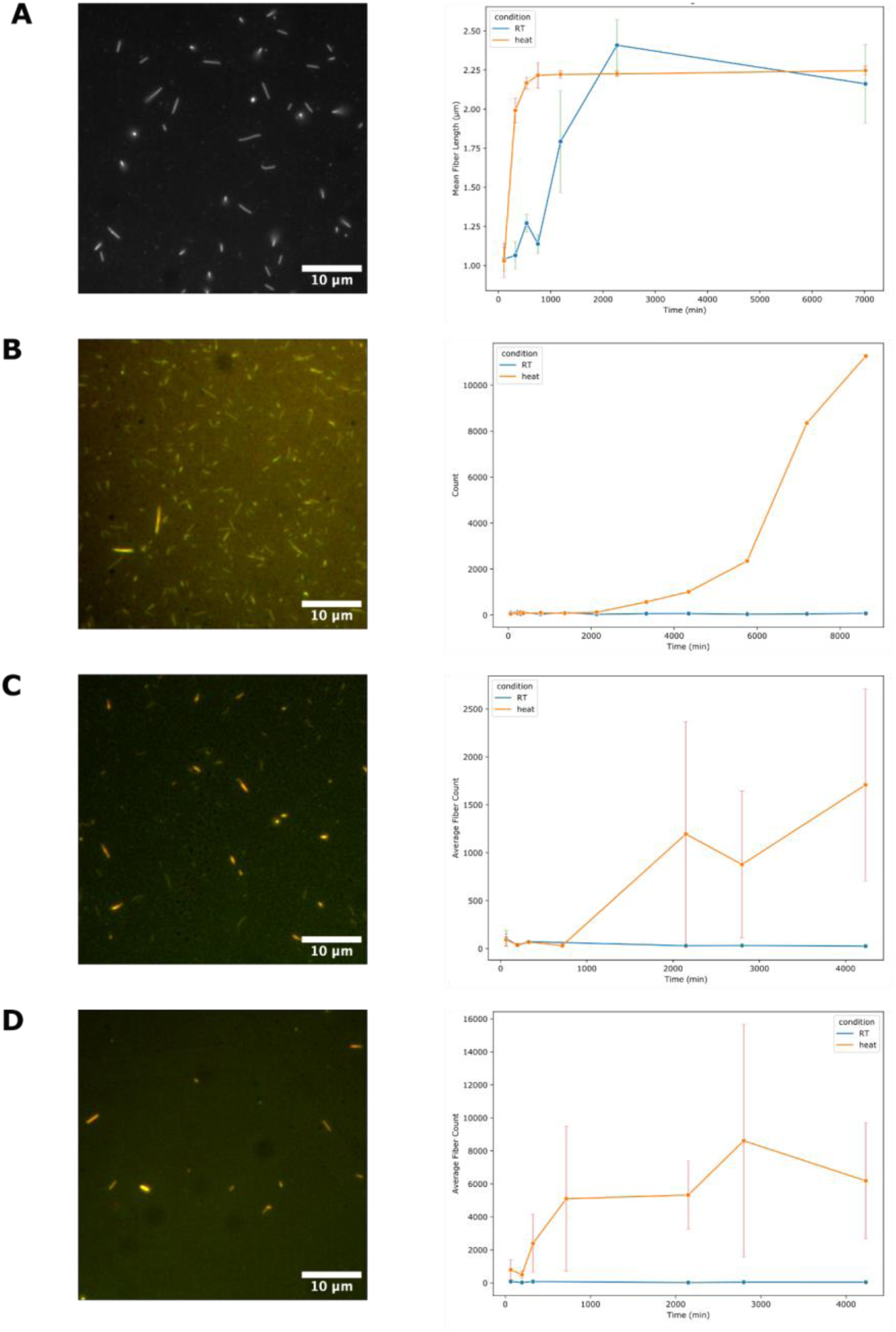
LHD_HF59 Fiber growth quantification of fluorescent protein variants and buffer conditions. (**A**) Quantification of average fiber length when both components are fused to GFP with representative fluorescent microscopy image on the left at 2130 minutes. (**B-C**) Quantification of fiber count for (**B**) B component fused to mScarlet-I with fluorescent image on the left at 7200 minutes. (**C**) B component fused to mScarlet-I with K to L mutation, fluorescent image on the left at 4350 minutes. (**D**) B component fused to mScarlet-I with K to L mutation kept in 400 mM Imidazole buffer, fluorescent image on the left at 4350 minutes.

**Fig. S6.**
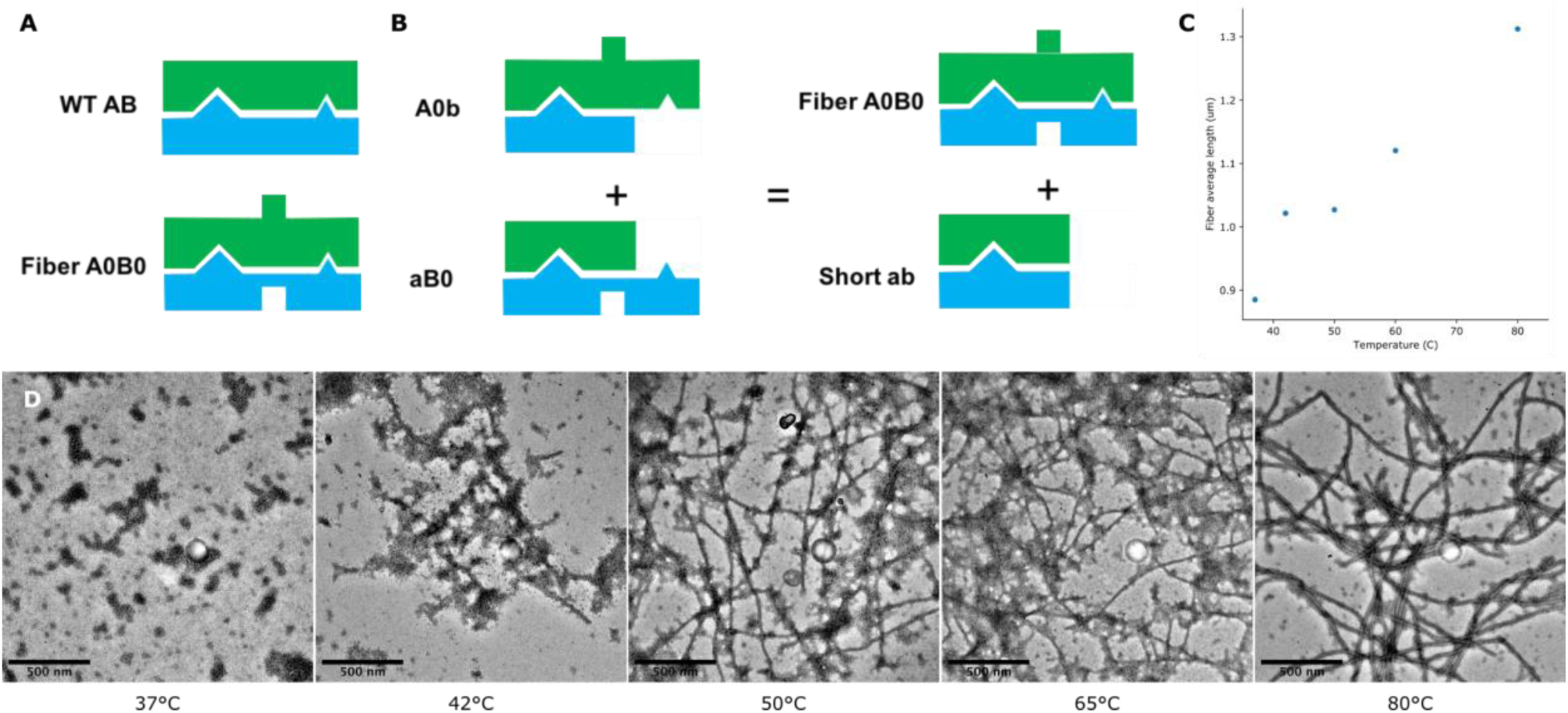
Fiber formation by mixing components coexpressed with temperature blocks. (**A**) Cartoon representation of wildtype (WT) non-fiber forming heterodimer scaffold AB (top) and fiber forming heterodimer (bottom) A0B0. (**B**) Cartoon representation of mixing A0 coexpressed with shorter complementary block b and B0 coexpressed with shorter complementary block a, and through thermodynamic exchange, the A0B0 and ab are formed. (**C**) Quantification of mixing A0 and B0 vs A0b and aB0 for 20 minutes at 37°C, 42°C, 50°C, 65°C, and 80°C by fluorescent microscopy and (**D**) representative electron microscopy images.

**Fig. S7.**
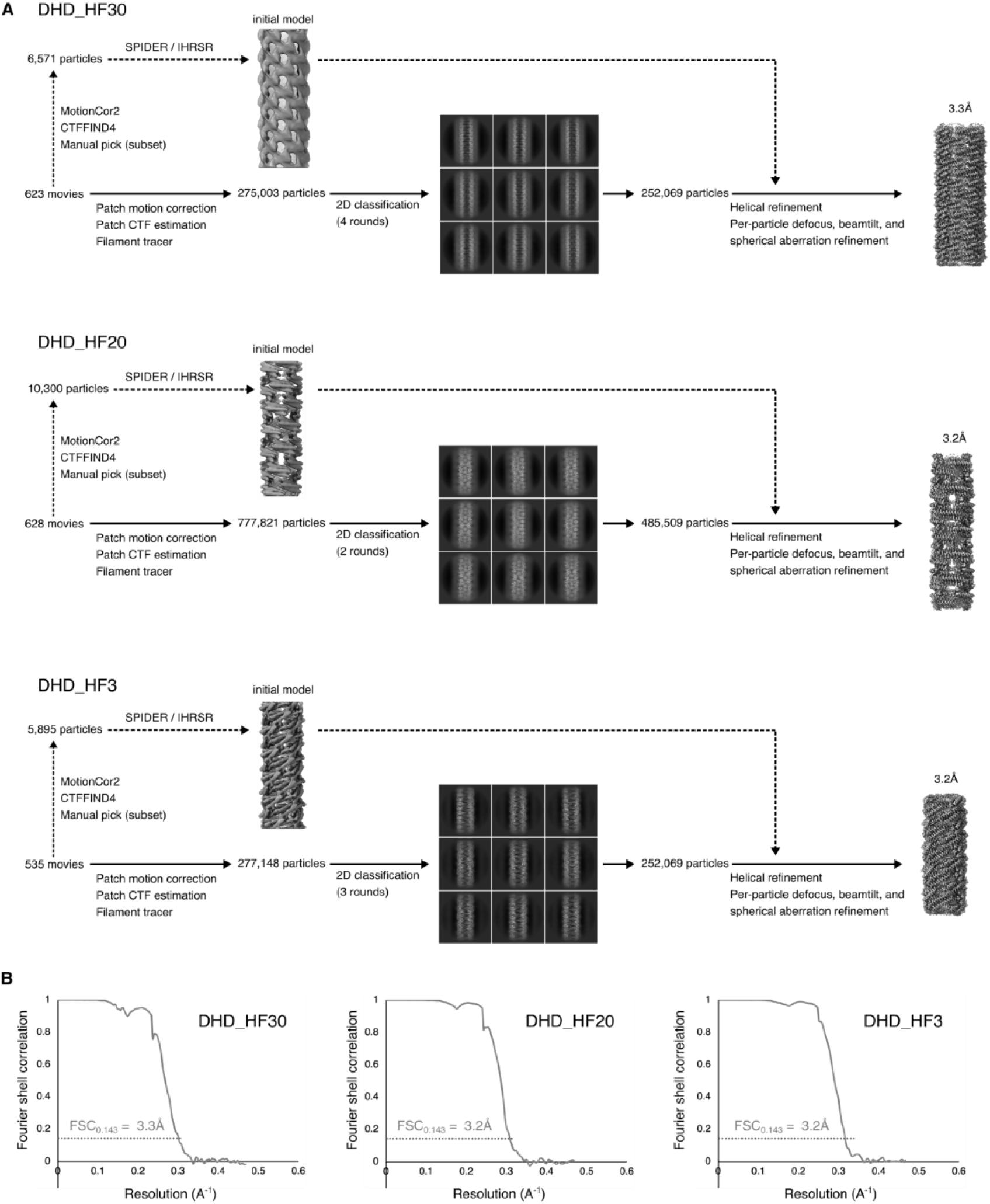
CryoEM data processing of two-component filaments. Flowcharts of cryoEM data processing **(A)** and Fourier shell correlation (FSC) curves **(B)** for DHD_HF30, DHD_HF20, and DHD_HF3.

**Fig. S8.**
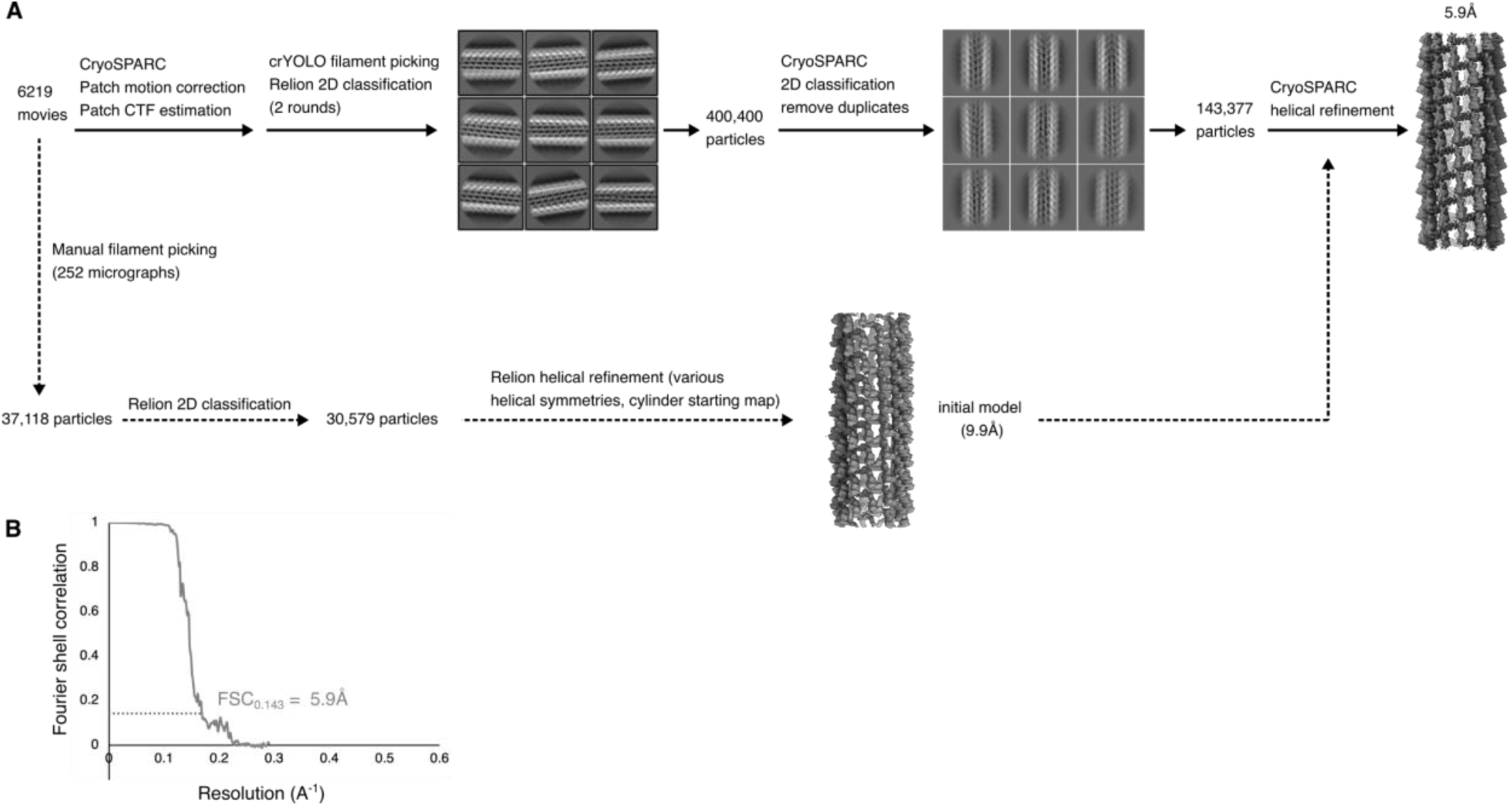
CryoEM data processing of LHD_HF59. Flowchart of cryoEM data processing **(A)** and Fourier shell correlation (FSC) curve **(B)** for LHD_HF59.

**Table S1:** Cryo-EM data collection, refinement, and validation statistics.

|  |  |  |  |  |
| --- | --- | --- | --- | --- |
| Structure | DHD_HF30 | DHD_HF20 | DHD_HF3 | LHD_HF59 |
| PDB code | 9NMK | 9NMJ | 9NMI | 9NML |
| EMDB code | EMD-49532 | EMD-49531 | EMD-49530 | EMD-49533 |
| Magnification | 130,000 | 130,000 | 130,000 | 45,000 |
| Voltage (kV) | 300 | 300 | 300 | 200 |
| Electron dose (e<sup>-</sup>/Å<sup>2</sup>) | 90 | 90 | 90 | 50 |
| Defocus range (μM) | 0.9-2.4 | 0.9-2.2 | 0.9-2.2 | 0.6-2.0 |
| Pixel size (data collection) (Å) | 0.525 | 0.525 | 0.525 | 0.885 |
| Pixel size (reconstruction) (Å) | 1.05 | 1.05 | 1.05 | 0.885 |
| Point group symmetry | C5 | C2 | C1 | C4 |
| Helical rise (Å) | 22.3 | 15.2 | 5.9 | 23.5 |
| Helical rotation (degrees) | 37.1 | 66.5 | -135.9 | 119.3 |
| Particle images | 86,025 | 485,509 | 252,069 | 146,028 |
| Resolution (FSC 0.143) (Å) | 3.3 | 3.2 | 3.2 | 5.9 |
| r.m.s. deviation bond lengths (Å) | 0.28 | 0.27 | 0.29 | 0.41 |
| r.m.s. deviation bond angles (degrees) | 0.61 | 0.55 | 0.61 | 0.59 |
| MolProbity score | 1.07 | 1.34 | 1.03 | 0.84 |
| Clashscore | 2.8 | 6 | 2 | 1.2 |
| C-beta deviations | 0 | 0 | 0 | 0 |
| Rotamer outliers (%) | 1 | 0 | 0 | 0 |
| Ramachandran favored (%) | 100 | 100 | 98 | 100 |
| Ramachandran allowed (%) | 0 | 0 | 2 | 0 |
| Ramachandran outliers (%) | 0 | 0 | 0 | 0 |

## Methods

### Computational Design Approach

The helical docking and design methodology^3^ was applied to multi-component proteins (DHD131, DHD37_1:234, DHD127, DHD15, DHT03, LHD29, LHD101, and LHD317) to create models of helical filament designs. The multi-component protein scaffolds were first treated as single-component scaffolds, but only those with fiber interface spanning multi-components were selected. The design trajectories were screened based on the following criteria: a Rosetta Energy Unit difference exceeding −15.0 between the polymeric (bound) and monomeric (unbound) states, an interface surface area greater than 700 Å², a Rosetta shape complementarity score above 0.62, and fewer than five unsatisfied polar residues. Designs that satisfied these thresholds were further refined manually, with non-essential mutations reverted to their native state. The highest-scoring design for each configuration was selected and included in the final protein set for experimental testing. Complete sequences are included in the supplementary information.

### Protein Expression and Purification

Synthetic genes were optimized for expression in *Escherichia coli* and synthesized by genscript and IDT. Bicistronic or tricistronic genes were cloned into pET29b+ vectors with either N- or C-terminal GFP fusions when space was adequate, using the NdeI and XhoI restriction sites for cloning. In these constructs, only one component contained a hexahistidine tag for purification via immobilized metal affinity chromatography (IMAC).

The constructs were transformed into BL21* (DE3) competent *E. coli* cells and grown in 50 mL of Terrific Broth medium containing 200 mg L⁻¹ kanamycin. Protein expression was induced using the Studier autoinduction method^14^ at 37°C for 24 hours. After induction, cells were harvested by centrifugation, and pellets were resuspended in Tris-buffered saline (TBS) and lysed with Bugbuster detergent. The soluble protein fraction was clarified by centrifugation and purified using Ni^2+^-based IMAC on Ni-NTA Superflow resin. During purification, the resin-bound lysate was washed with 10 column volumes of buffer containing 40 mM imidazole and 500 mM NaCl, followed by elution with buffer containing 400 mM imidazole and 75 mM NaCl. Both soluble and insoluble fractions were analyzed by SDS-PAGE, and samples with protein bands at the expected molecular weight were selected for electron microscopy screening. For further characterization, selected designs were scaled up to 0.5 L cultures, expressed under identical conditions for 24 hours at 37°C, and harvested by centrifugation. Cell pellets were resuspended in TBS, lysed via microfluidization, and purified following the same method.

### Negative Stain Electron Microscopy

For electron microscopy screening, the soluble fractions were concentrated in a buffer containing 25 mM Tris, 75 mM NaCl, and adjusted to pH 8. A 6 µL droplet of the sample, prepared by diluting 1 µL of protein with 5 µL of buffer, was applied to carbon-coated 200-mesh copper grids that had been negatively glow-discharged. The grids were then rinsed with Milli-Q water and stained with 0.75% uranyl formate (pH 4.0) following established protocols^15^. Screening was conducted using a 120 kV Talos L120C transmission electron microscope (ThermoFisher). Images were acquired with a bottom-mounted Teitz CMOS 4k camera system and further processed to enhance contrast using Fiji software^16^.

### Cryo-Electron Microscopy

CryoEM samples were prepared by adding protein to glow-discharged CFLAT 2/2 holy-carbon grids, blotting away liquid, and plunging into liquid ethane using a Vitrobot (ThermoFisher). For DHD_HF30, DHD_HF20, and DHD_HF3, movies were acquired on a Krios microscope (ThermoFisher) equipped with a K-2 Summit Direct Detect camera (Gatan Inc.) operating in superresolution mode, with a pixel size of 0.525 Å/pixel, 50 frames, and a total dose of 90 electrons/Å^2^. Data collection was automated using Leginon^17^. The data processing workflow is summarized in Fig. S7. Movies were aligned, dose-weighted, and binned to 1.05 Å/pixel using MotionCor2^18^, and CTF parameters were estimated using CTFFIND4^19^. Helices were manually picked from a subset of images using Appion. Helical symmetry parameters and an initial structure were determined using IHRSR^8,9^ in SPIDER. Further processing was then performed in cryoSPARC^12^. Movies were aligned, dose-weighted, and binned to 1.05 Å/pixel using patch motion correction, and CTF parameters were estimated by patch CTF. Helices were picked using the filament tracer, then subjected to multiple rounds of 2D classification. Particles from high-quality 2D classes were then subjected to helical refinement, using the starting helical symmetry parameters and maps obtained from IHRSR in SPIDER. Per-particle defocus, beamtilt, and spherical aberration were also refined. For LHD_HF59, movies were acquired on a Glacios microscope (ThermoFisher) equipped with a K3 Direct Detect camera (Gatan Inc.) operating in counting mode, with a pixel size of 0.885 Å/pixel, 99 frames, and a total dose of 50 electron/Å^2^. Data collection was automated with SerialEM^20^. The data processing workflow is summarized in Fig. S8. Movies were aligned and dose-weighted using patch motion correction, and CTF parameters were estimated by patch CTF using cryoSPARC. Filaments were picked manually from a subset of micrographs using the Relion^10,11^ helical picker, then subjected to Relion 2D classification. Particles belonging to high-quality 2D classes were selected for 3D refinement in Relion. Helical symmetry parameters and an initial structure were determined by performing a series of refinements with various helical symmetry parameters and a cylinder starting model. Filaments were picked automatically from the full dataset using crYOLO^21,22^, following training with the manually picked particles, then subjected to multiple rounds of 2D classification in Relion and cryoSPARC. A final helical reconstruction was performed in cryoSPARC, using the starting model and helical symmetry parameters determined using Relion. Atomic models were built using ISOLDE^23^ and real-space refinement in Phenix^24^. Cryo-EM data collection, refinement, and validation statistics are summarized in Table S1.

### Fluorescence Microscopy

#### Sample Preparation

To prepare samples for imaging, 1 µL of the fiber solution was diluted in 1 mL of TBS buffer (25 mM Tris, 75 mM NaCl, pH 8). From this dilution, 50 µL was added to a well for imaging.

#### Image Acquisition

Fiber samples were immobilized directly onto the glass surface of a 384-well plate (Greiner Bio-One, cat# 781892, https://www.gbo.com/), allowed to adhere for 5 minutes, and subsequently washed to minimize background signal. Images were captured using an IN Cell Analyzer 2500HS equipped with a 60X 0.95 NA Plan Apo CFI objective lens (Nikon, 12605). Fiber fluorescence was excited using a seven-color Solid State Illuminator (SSI), and sequential fluorescent signals were acquired with the following filter settings: Green (excitation 473/28 nm, emission 511/23 nm) and Red (excitation 575/30 nm, emission 623/42 nm). Imaging was managed using IN Cell Analyzer 2500HS software version 7.4, and data were collected on an sCMOS camera without binning. For each condition, 15–25 fields of view, spaced evenly across the well, were imaged.

### Image Analysis

Fiber images were segmented using a custom pipeline developed in a Jupyter notebook. A subset of 35 images was reserved for training a random forest classifier. These training images were manually annotated to identify true positive fiber pixels, which were then used to train the classifier utilizing the RandomForestClassifier module from the Scikit-learn library^25^. The classifier was configured with a maximum tree depth of 10 and 50 estimators. The training achieved accuracy scores exceeding 0.99, and the segmentation results were qualitatively comparable to a previous model created using the Weka Trainable Segmentation plugin in Fiji^16,26^.

Once the model was trained, a separate analysis notebook was employed to perform segmentation on all images. Post-segmentation, small objects (fewer than 64 pixels) were filtered out, and a binary closing operation was applied to address potential over-segmentation. Fiber counts were determined by the number of identified fiber objects, and fiber lengths were measured based on the major axis length of each object.

### Atomic Force Microscopy

AFM imaging was conducted in liquid using AC tapping mode at 25 °C on a Cypher ES (Asylum Research). A Nanoworld probe (k = 0.15 N/m, f = 1200 kHz) was used, with a scanning rate of 1.5 Hz (512 lines per image) and a drive frequency of ∼500 kHz. The amplitude setpoint was adjusted to preserve surface topography. Data analysis was performed using Gwyddion. Imaging was done in a 70–100 µL droplet of 1 µM protein solution on a mica surface.

### Cell lines

Cell culture experiments were performed in HEK293t cells (ATCC #CRL 3216). Cell cultures were tested monthly for mycoplasma contamination (Southern Biotech #13100–01).

### In-vivo cell culture experiments

The open reading frame of B0 and AS were cloned into the pCDNA 3.1 (+) vector backbone; similarly, the open reading frame of A0 was cloned into the same vector backbone with a C-terminal eGFP tag. HEK293t cells were plated at 2×10^5^ cells /ml in 10% FBS DMEM on a glass-bottom 24-well plate. Twenty-four hours after plating, cells were transfected with 500 ng BS plasmid, as well as 250 ng of both the B0 and A0 plasmids. Twenty-four hours following transfection, cells in the starvation group were starved by replacing media with 0% FBS DMEM media. Similarly, cells receiving drug treatment were dosed with 20 mM 8-Br-cAMP. Twenty-four hours after drug treatment and starvation, cells were imaged using a CSU-X1 spinning disk confocal microscope. Z-stack images of the GFP channel were collected at 40X magnification with an additional 1.7x Gataca superresolution magnification module. In triplicate, Z-stacks were taken with a 0.6 um spacing between slices, compiled, and analyzed for comparison across treatment groups. Uncropped Z-stack images were used to quantify the frequency of the fiber-forming phenotype in cells across treatment groups. Cells were considered to exhibit fiber formation if the cell contained a strand longer than 5μm in length. Cells without this categorization, or without any discernible fibers, were considered to not exhibit fiber formation.

### Data availability

Cryo-EM maps and atomic models have been deposited in the Electron Microscopy Data Bank (EMDB) and Protein Data Bank (PDB), respectively, with the following accession codes: EMD-49530, PDB: 9NMI (DHD_HF3 Filament); EMD-49531, PDB: 9NMJ (DHD_HF20 Filament); EMD-49532, PDB: 9NMK (DHD_HF30 Filament); EMD-49533, PDB: 9NML (LHD_HF59 Filament).

